# Rcompadre and Rage - two R packages to facilitate the use of the COMPADRE and COMADRE databases and calculation of life history traits from matrix population models

**DOI:** 10.1101/2021.04.26.441330

**Authors:** Owen R. Jones, Patrick Barks, Iain Stott, Tamora D. James, Sam Levin, William K. Petry, Pol Capdevila, Judy Che-Castaldo, John Jackson, Gesa Römer, Caroline Schuette, Chelsea C. Thomas, Roberto Salguero-Gómez

## Abstract

1. Matrix population models (MPMs) are an important tool for biologists seeking to understand the causes and consequences of variation in vital rates (e.g., survival, reproduction) across life cycles. Empirical MPMs describe the age- or stage-structured demography of organisms and usually represent the life history of a population during a particular time frame at a specific geographic location.
2. The COMPADRE Plant Matrix Database and COMADRE Animal Matrix Database are the most extensive resources for MPM data, collectively containing >12,000 individual projection matrices for >1,100 species globally. Although these databases represent an unparalleled resource for researchers, land managers, and educators, the current computational tools available to answer questions with MPMs impose significant barriers to potential COM(P)ADRE database users by requiring advanced knowledge to handle diverse data structures and program custom analysis functions.
3. To close this knowledge gap, we present two interrelated R packages designed to (i) facilitate the use of these databases by providing functions to acquire, quality control, and manage both the MPM data contained in COMPADRE and COMADRE, and a user’s own MPM data (**`Rcompadre`**), and (ii) present a range of functions to calculate life history traits from MPMs in support of ecological and evolutionary analyses (**`Rage`**). We provide examples to illustrate the use of both.
4. **`Rcompadre`** and **`Rage`** will facilitate demographic analyses using MPM data and contribute to the improved replicability of studies using these data. We hope that this new functionality will allow researchers, land managers, and educators to unlock the potential behind the thousands of MPMs and ancillary metadata stored in the COMPADRE and COMADRE matrix databases, and in their own MPM data.

## Introduction

Matrix population models (MPMs, hereafter) have become a commonplace tool for ecologists, evolutionary biologists, and conservation biologists seeking to understand how variation in vital rates (e.g., survival, development, reproduction, recruitment, etc.) in the life cycle varies geographically and across species. MPMs describe population dynamics based on stage- or age-specific vital rates in the population of interest over their life cycle (Caswell, 2001). Outputs derived from MPMs include population growth rates (Caswell, 2001), key life-history traits (Caswell, 2001), and vital rate sensitivities (de Kroon, Plaisier, van Groenendael, & Caswell, 1986; de Kroon, van Groenendael, & Ehrlén, 2000). These outputs each have a well-understood biological interpretation, which allows comparison of MPM-derived population and life history metrics, and thus demography across the diversity of life on Earth, from moss (e.g., Okland, 1995) to monkeys (e.g., Morris et al., 2011) to microbes (e.g., Jouvet, Rodríguez-Rojas, & Steiner, 2018), and in myriad ecoregions.

Since the introduction of MPMs in the 1940s (Leslie, 1945, 1948), researchers have published thousands of MPMs for thousands of species. Our team has been digitising these MPMs into centralised databases for plants (the COMPADRE Plant Matrix Database: Salguero-Gómez et al., 2015) and animals (the COMADRE Animal Matrix Database: Salguero-Gómez et al., 2016). These twin databases now contain more than 12,000 MPMs for more than 1,100 species (COMPADRE: 8,708 matrices for 757 species; COMADRE: 3,317 matrices for 415 species, as of September 2021) and are regularly augmented with newly-published and newly-digitised records. The databases, their history, and the rationale behind the data organisation are described in Salguero-Gómez et al. (2015) and Salguero-Gómez et al. (2016), respectively.

COMPADRE and COMADRE store and provide MPMs and their associated metadata in a hierarchical structure that, while efficient for distribution, can be both a barrier to use and an entry point for user errors. The primary component of MPMs are the two-dimensional, square projection matrices, and the size of these matrices can vary widely across species and studies. Moreover, most projection matrices (**A**) in the databases are partitioned into their three constituent process-based submatrices such that **A** = **U** + **F** + **C**. Here, submatrix **U** describes transitions related to survival and growth/development, submatrix **F** describes sexual reproduction, and submatrix **C** describes clonal reproduction. Thus, in most cases, each MPM is represented by these four matrices (**A**, the main projection matrix and the submatrices **U**, **F** and **C**) alongside information about the life cycle stages used in the MPM. In the majority of cases, the projection interval (time step) for the MPM is one year, but this can vary considerably depending on the life history of the organism concerned (for example, five year intervals are common in tree MPMs). Each MPM in the databases is also associated with over 40 metadata variables extracted from its parent original work(s) (e.g., stage definitions, projection time steps, citation, taxonomy, geography, etc., detailed in Salguero-Gómez et al., 2015 & 2016). This nested structure allows for higher digitisation fidelity and distribution efficiency, but also means that the dataset cannot be imported by ordinary spreadsheet software, such as Excel, which accommodate only rectangular (or “flat”) data structures. Both of the most common tools for working with MPMs, the R statistical programming language (R Core Team, 2021) and Matlab (Matlab, 2010), readily accept hierarchical data structures. However, users must have a familiarity with handling a range of nested object classes to organise the databases to suit their needs (e.g., “*subset to only primates*” or “*subset to only species from tropical ecoregions*”). The higher dimensionality can increase the risk of errors, such as using the wrong data dimension, even for experienced users.

The R package ecosystem provides a wide range of tools for analysing population dynamics from MPMs within individual populations. For example, **`popdemo`** (Stott, Hodgson, & Townley, 2012) focuses on the calculation of metrics related to transient population dynamics and transfer function analyses; **`popbio`** (Stubben, Milligan, & Others, 2007) provides functions to accomplish many (but not all) of the analyses found in the textbooks of Caswell (2001) and Morris & Doak (2002), such as the calculation of eigen properties (i.e., the asymptotic population growth rate, stable stage structure and reproductive values) or sensitivities and elasticities; **`Rramas`** (de la Cruz Rot, 2019) provides tools for making population projections and conducting population viability analyses from MPM data; and **`lefko3`** (Shefferson, Kurokawa, & Ehrlén, 2021) provides tools that allow the inclusion of information on individual histories, which could influence population dynamics, into MPM analyses (see Ehrlén, 2000). However, the tools for life history analysis provided by these existing packages are more limited, with among the most notable absence being important life history metrics based on age-from-stage calculations. Researchers that wanted to make such calculations (e.g., measures of senescence, longevity, or age at maturity) have needed to write their own code based on published equations in mathematics-heavy work, which has been a barrier to the broader adoption of these methods. Moreover, these life history metrics are often most meaningful in analyses across many populations or species. The existing packages provide little support for the large hierarchical data structures needed to apply analyses to hundreds or thousands of MPMs that may underlie a single comparative or macroecological analysis.

Here, we introduce two R packages that enable users to construct robust MPM analysis workflows to answer questions from single populations to across the tree of life. The first package, **`Rcompadre`**, is designed to facilitate acquisition, quality control, and management of the rich, hierarchical MPM data in COMPADRE and COMADRE. For example, this package includes tools to filter (subset) the databases based on metadata archived in these resources (e.g., by ecoregion, by taxonomic group). In addition to “base” style R syntax for these tasks, **`Rcompadre`** integrates **`tidyverse`** (Wickham et al., 2019) functionality to improve usability. The second package, **`Rage`**, builds on the enhanced data accessibility provided by **`Rcompadre`** by providing analysis pipeline support for arbitrarily large numbers of MPMs and the calculation of life history traits needed to support comparative analyses on this scale. These life history traits include life tables, mean life expectancy, generation time, among several others.

We showcase downloading, subsetting, and preparing MPM data for a broad comparative analysis using publicly-accessible data retrieved with **`Rcompadre`** (Box 1). We then illustrate an application of **`Rage`** to calculate ecologically and evolutionarily relevant metrics to test hypotheses related to life history theory at broad taxonomic scale. In doing so, we demonstrate the functional integration of **`Rcompadre`** and **`Rage`** and how investigators can use them in tandem to design workflows (Fig. 1) to answer their own questions in ecology, evolution and conservation biology.

**Figure 1.**
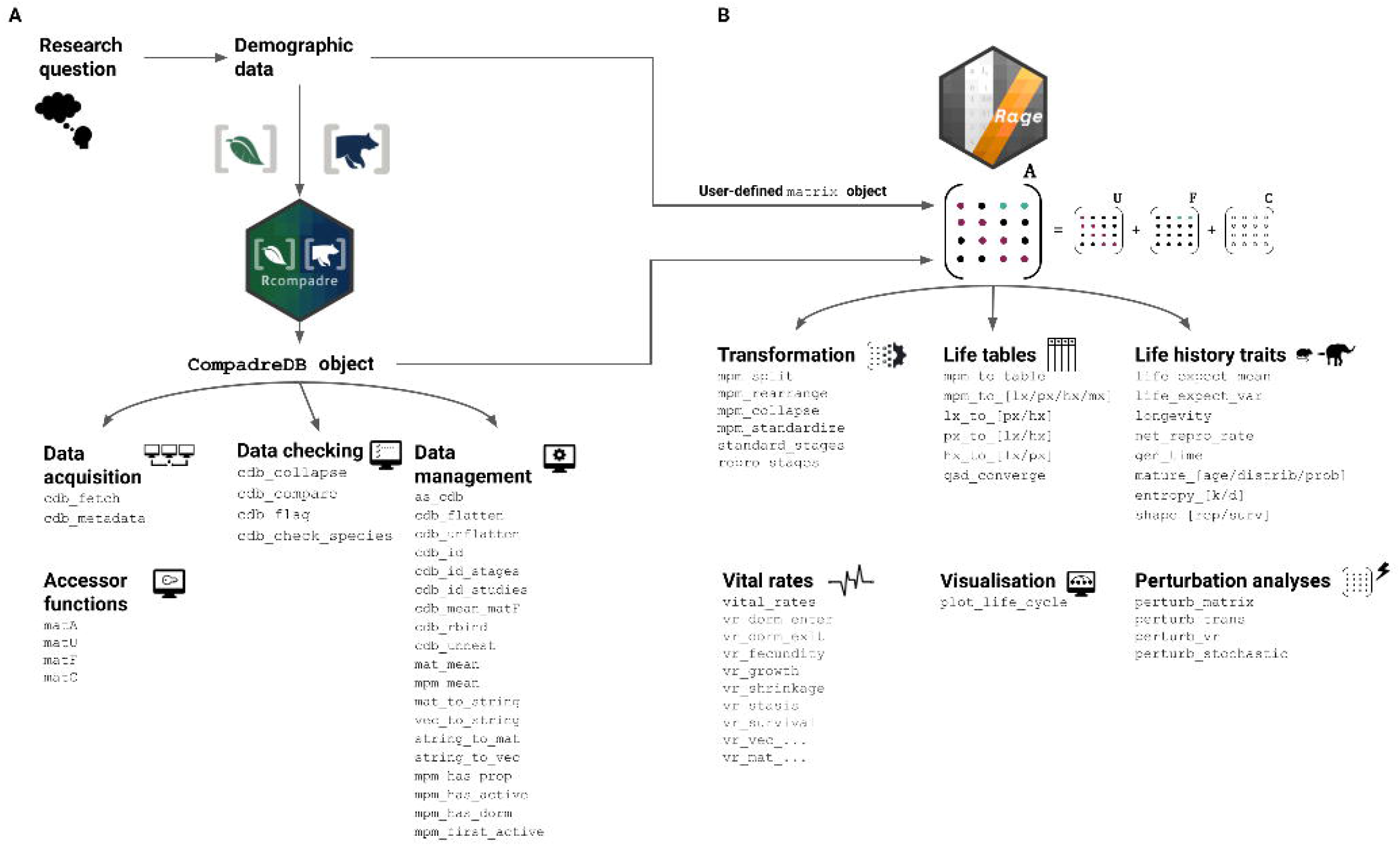
Workflow of using **`Rcompadre`** and **`Rage`** for ecological and evolutionary analyses of matrix population model data. (**A**) Once the author(s) have identified the research question, demographic data in the format of MPMs can be accessed from the COMPADRE and/or COMADRE databases via the **`Rcompadre`** R package. This package allows for the online acquisition, checking (according to data needs) and management of the `CompadreDB` data object (e.g., using `cdb_fetch` to download the data and `cdb_flag` and `filter/subset` to produce a data set for analysis). (**B**) The filtered data (or other user-provided MPM data) can be then migrated for calculations of life history traits with **`Rage`** (alternatively, these can be done directly on MPMs provided by the author). The families of functions archived in **`Rage`** include: transformation (e.g., `mpm_collapse`), creation of life tables (e.g., `mpm_to_lx`), derivation of life history traits (e.g., `longevity`), calculation of vital rates (e.g., using `vital_rates` to calculate average survival, reproduction, development, etc.), visualisation of life cycles (e.g., `plot_life_cycle`), and perturbation analyses (e.g., `perturb_stochastic`).

**Figure 2.**
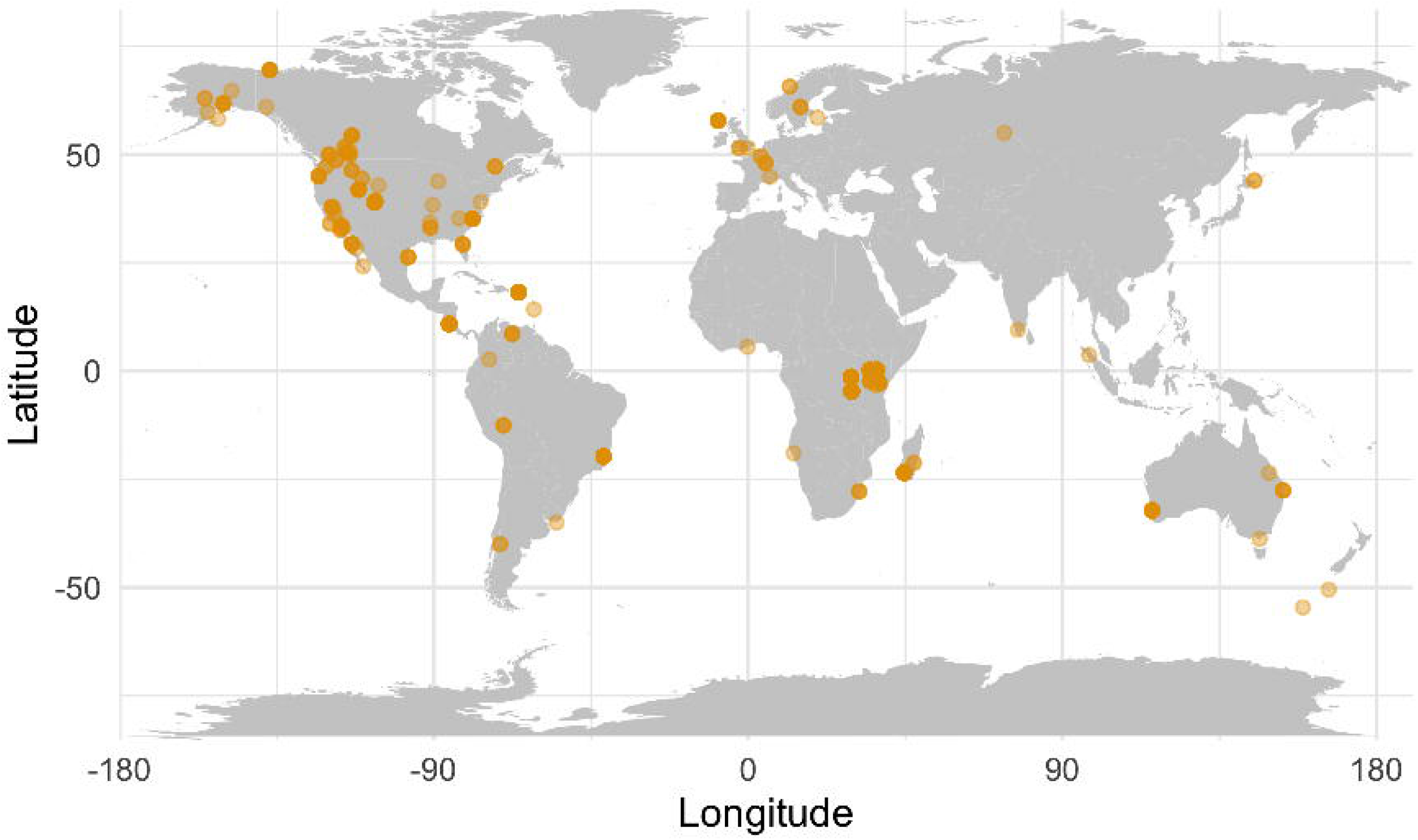
The spatial extent of data in the subset of mammal data used in our example analysis. Note that 186 of the matrices for mammals in our set (~27%) lack associated spatial information.

**Figure 3.**
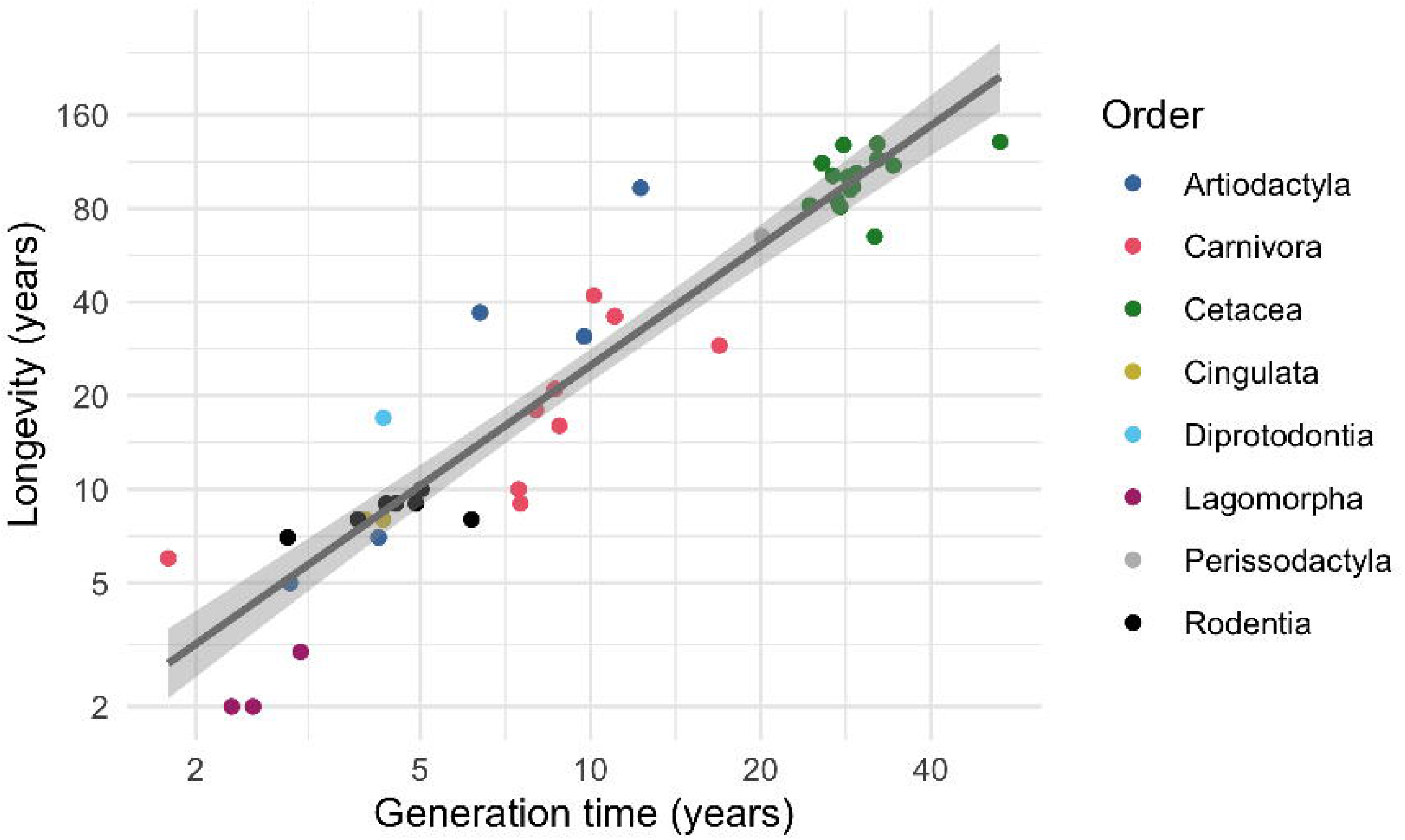
The relationship between estimated generation time and longevity (defined as the age that 1% of a synthetic cohort would reach, based on the MPM). The line represents the fit of an ordinary least-squared regression through the data. The slope is 1.28 (±0.07) and the intercept is 0.26 (±0.16); R^2^=0.90; F_1,43_= 379; p <0.001).

### Box 1: Using Rcompadre to download and prepare MPM data for analysis

In the following example, we illustrate the use of `Rcompadre` to carry out typical data download and preparation tasks for an analysis relevant to comparative population dynamics research. Specifically, we aim towards an analysis of mammalian life span and its relationship with generation time (continued in **Box 2**). After loading the required packages, we download the COMADRE data and conduct some basic checks of the matrices. We then filter the data set to include only mammals, to include no missing values in the **U** matrix, and to ensure that the **U** and **F** matrices are not filled entirely with zero values, nor that columns of the **U** matrix sum to 0. We further filter the data to ensure that the projection interval is 1 year. Finally, we can plot the geographic distribution of these data using tools from the `ggplot2` and `maps` packages (Fig. 2).

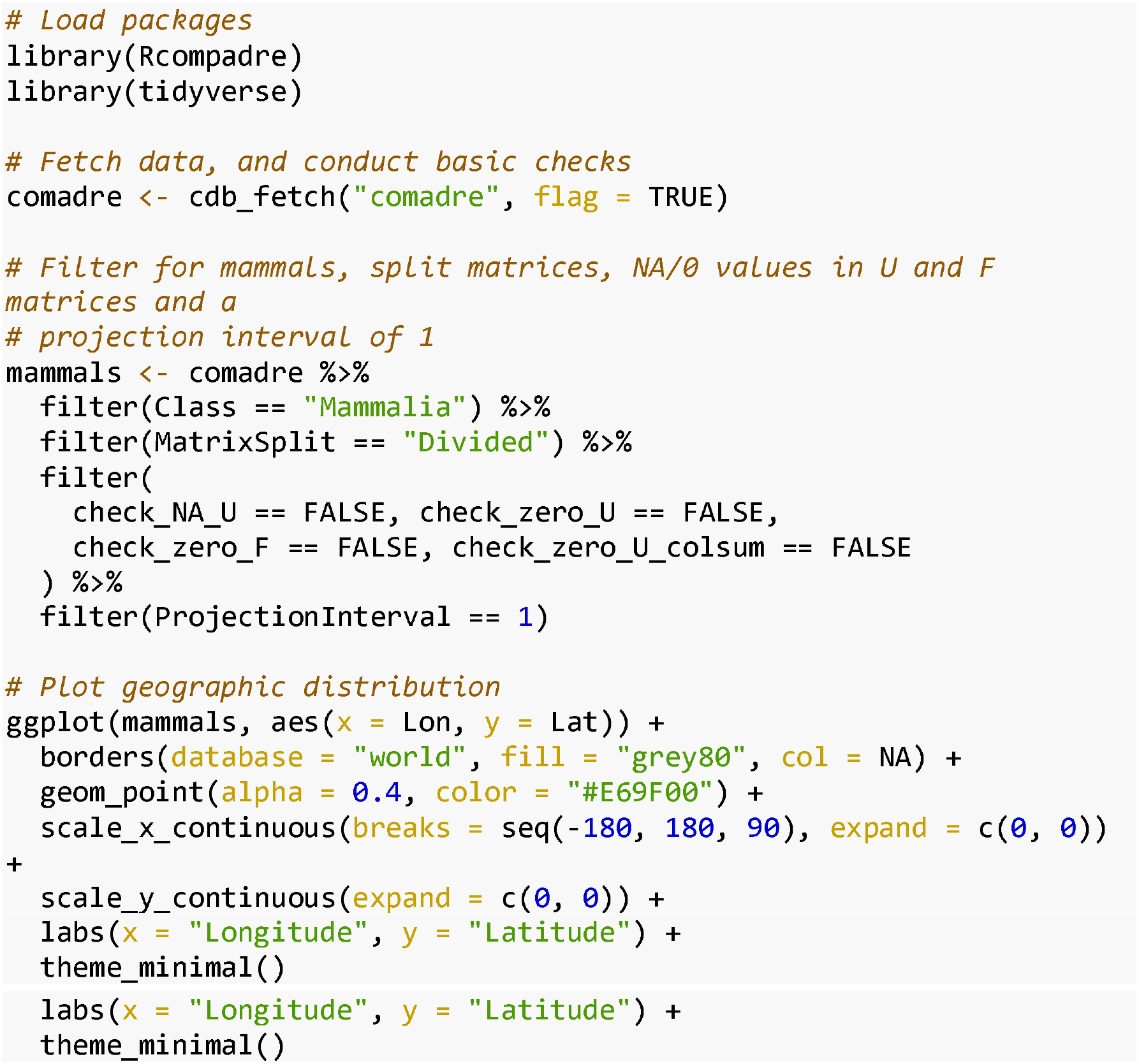

## Rcompadre

**`Rcompadre`** contains functions to facilitate downloading and using MPMs alongside their metadata from the COMPADRE and COMADRE databases (Fig. 1a). A central feature of this package is the definition of a new object class, `CompadreDB`, which allows R functions that are already familiar to users (e.g., `head` or **`tidyverse`** verbs) to be augmented with ‘methods’ that ensure that they appropriately handle the structure of MPM data from the COM(P)ADRE databases. In addition to improving user-friendliness, the class definition provides a pathway for extending the compatibility of COM(P)ADRE data to other existing or future R functions. Briefly, the structure of `CompadreDB` objects uses the S4 systems^1^ with two slots: (1) the `data` slot, which contains a `tibble`-style data frame (Wickham & Grolemund, 2016) with a list-column of MPMs and vector columns of metadata, and (2) the `version` slot which contains database version information for reproducibility, including the version number, date created, and a link to the database user agreement. In addition, we have created the `CompadreMat` class, which formally defines how MPMs are represented in a `CompadreDB` object. Here too, the use of an explicit class definition has allowed us to define how the data contained in the object will respond to familiar R functions. For example, users can access and replace columns of data using the standard `x$name` and `x$name <-value` methods, respectively. In addition, we provide the functionality to access the matrix data directly, for example, using the functions `matA` or `matU` to access all **A** matrices or **U** submatrices in the database as a list. This functionality is particularly convenient if the user wishes to apply functions to a large set of MPMs, as one would do in comparative and macroecological analysis (for example, see recent studies by Coutts et al. (2016), Takada & Kawai (2020), James et al. (2020), Healy et al. (2019), Capdevila et al. (2020) and Jones et al. (2020)). In addition to ‘base’ R functions, many data analysis workflows make use of functions in the **`tidyverse`** family of packages (Wickham et al., 2019). Our package includes “tidy” methods for `CompadreDB` objects, allowing users to `filter, arrange, mutate, select, summarise, rename` and `join` COM(P)ADRE data to answer their study questions efficiently and at scale. The provision of these **`tidyverse`** methods also means that **`Rcompadre`** benefits from the piping (e.g., %>%) functionality of **`magrittr`** and more recently in base R (| >, in v.4.1.0 and later). Examples of how this functionality can streamline the human readability of workflows can be found in the vignettes at the package development pages.

In addition to a wide range of method-based support of existing R functions, **`Rcompadre`** provides functions for additional workflow tasks that follow the naming pattern of `cdb_` (pronounced “compadre database”) followed by a meaningful verb. For example, `cdb_fetch` retrieves COM(P)ADRE data of the current or any previous database version from the web as a `CompadreDB` object, and `cdb_compare` reports the differences between any pair of `CompadreDB` objects. Table 1 summarises the most important **`Rcompadre`** functions, and full documentation of all functions is provided in the package manual.

**Table 1.**
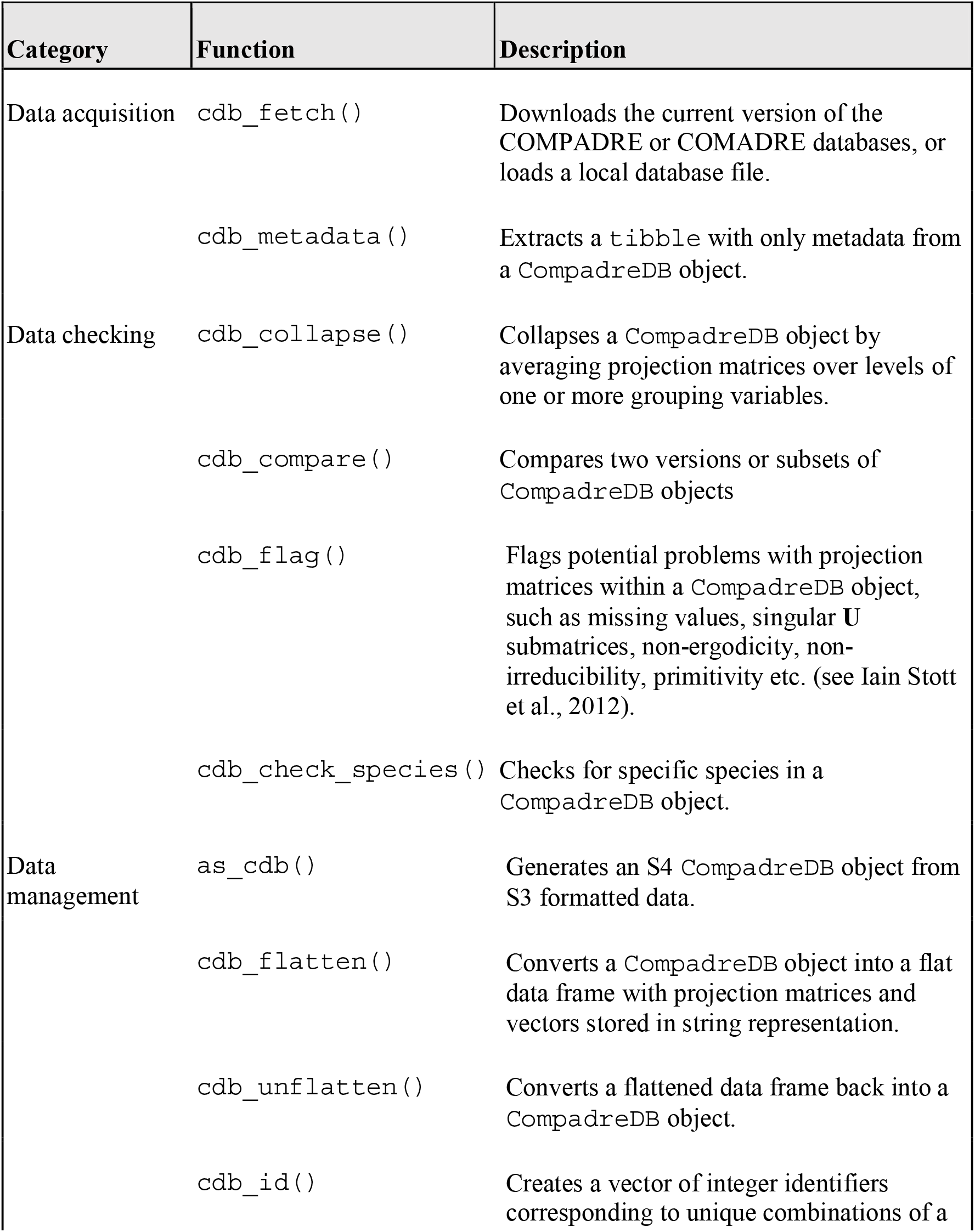
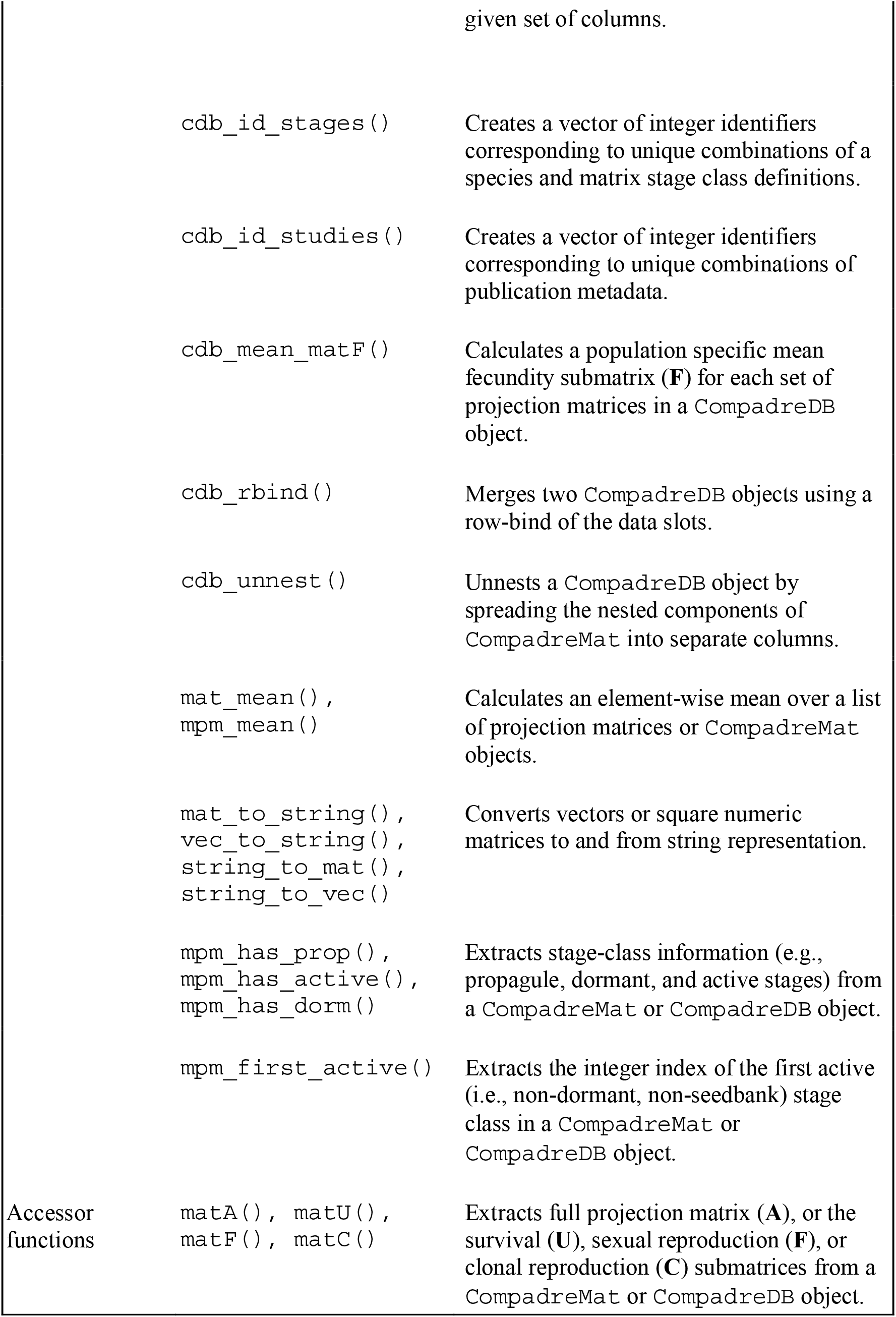
The functions in Rcompadre are grouped into four categories: Data acquisition, Data checking, Data management and Accessor functions. We outline the most important functions here, with a brief description. Users should consult the package documentation for a full description of named functions (e.g., `?cdb_fetch`) and to see a full list of functions.

### Data management and checking

The COM(P)ADRE databases include metadata associated with each MPM including taxonomic information, geolocation, and details of the source publication (see the User Guide at www.compadre-db.org or Salguero-Gomez et al. 2015, 2016 for full metadata documentation). When working with these data via Rcompadre, we can see the richness of the metadata with R’s `names` function and users can use any of these metadata columns to filter the database prior to analysis. The projection matrices themselves are contained in a list column called `mat`, where each element includes a list of the four matrices: **A** and the submatrices **U**, **F** and **C** (see above). The list also provides information on matrix stage definitions. All other columns of the COMADRE database object are ordinary vectors.

Not all COM(P)ADRE data will meet the inclusion criteria for a particular analysis. **`Rcompadre`** includes several general functions for checking the data that use the quality control flags generated when MPMs are digitised and checked before addition to the databases. These data checks are accessed through **`Rcompadre`** using the `cdb_flag` function. This function, which can be implemented as a stand-alone function or during data retrieval by `cdb_fetch`, adds logical metadata columns to the provided `CompadreDB` object which can be used for data filtering (see `?cdb_flag` for details of the available data property checks). For example, a minority of studies published only the main projection matrix, **A**, thereby preventing its decomposition into the **U**, **F** and **C** submatrices which may preclude certain demographic analyses. Matrices may also have missing (NA) values where a transition was not estimated. Other potential pitfalls flagged by this function include matrices that are singular (non-invertible), non-ergodic (where initial stage structure can influence asymptotic population growth rate), reducible (where the associated life cycle graph does not contain all necessary transition rates to enable pathways from all stages to all other stages) or non-primitive (Caswell, 2001; Stott, Townley, & Carslake, 2010). Depending on the desired downstream analyses, researchers may need to `filter` the database based on one or more of these flag columns.

The quality checks performed by `cdb_flag` cannot anticipate all potential inclusion criteria, and we strongly encourage investigators to perform additional checks that may be necessary to determine the suitability of a MPM record for their analysis. The existing metadata columns associated with each MPM contains a wealth of useful information to this end. For example, the interpretation of many metrics derived from MPMs depends on the projection interval (`ProjectionInterval`). We advise users to `filter` on this column to a common projection interval prior to analysis or to correct analysis outputs to the same temporal units. An analysis may also require delineating MPM records that use post-vs. pre-reproductive census models. Although both databases have a metadata field that reports this information (`CensusType`), it is often not reported in original publications and thus COM(P)ADRE includes records with incomplete metadata. Users may therefore need to carefully consider the source publication (e.g., retrieved using the `DOI_ISBN` and `AdditionalSource` column metadata) or contact the original authors to determine suitability.

Finally, **`Rcompadre`** includes a function, `cdb_build_cdb`, which allows users to access the full functionality of **`Rcompadre`** for their own data by constructing valid `CompadreDB` objects from user-supplied lists of matrices, (optional) stage information, and an accompanying data frame of metadata. Furthermore, we provide a way for users to augment COM(P)ADRE with a `CompadreDB` object containing their own data using the function `cdb_rbind`. This nimble data extensibility ensures the continued utility of **`Rcompadre`**’s suite of workflow tools without dependency on externally-maintained data.

In **Box 1** we illustrate the use of **`Rcompadre`** to download, check, and filter the COMADRE database (animal MPMs) in preparation for a later analysis of mammal life span using **`Rage`**. Vignettes at the **`Rcompadre`** documentation website (https://jonesor.github.io/Rcompadre/) give further detailed coverage of the package’s capabilities.

### Rage

The **`Rage`** package contains functions to facilitate the calculation of life history metrics (Table 2) from MPMs. The guiding philosophy of the package centres on (i) augmenting the suite of life history analyses that are implemented in R and (ii) providing support for analyses—whether new in **`Rage`** or previously implemented elsewhere—to be conducted in a standardised way across large numbers of MPMs. Other functions are novel, such as estimates of the pace and shape of reproduction (Baudisch & Stott, 2019). Broadly, the functions fall into six categories (Fig 1B, Table 2):

1. Transformation: reshape, resize, and reorder whole MPMs
2. Life tables: convert MPMs to life tables and life table components
3. Life history traits: calculate life history metrics
4. Vital rates: extract and summarise the component vital rates of MPMs
5. Visualisation: plot the life cycle graph
6. Perturbation analyses: calculate sensitivity and (stochastic) elasticity of any demographic statistic to perturbations of MPM elements, vital rates, or transition types

**Table 2.**
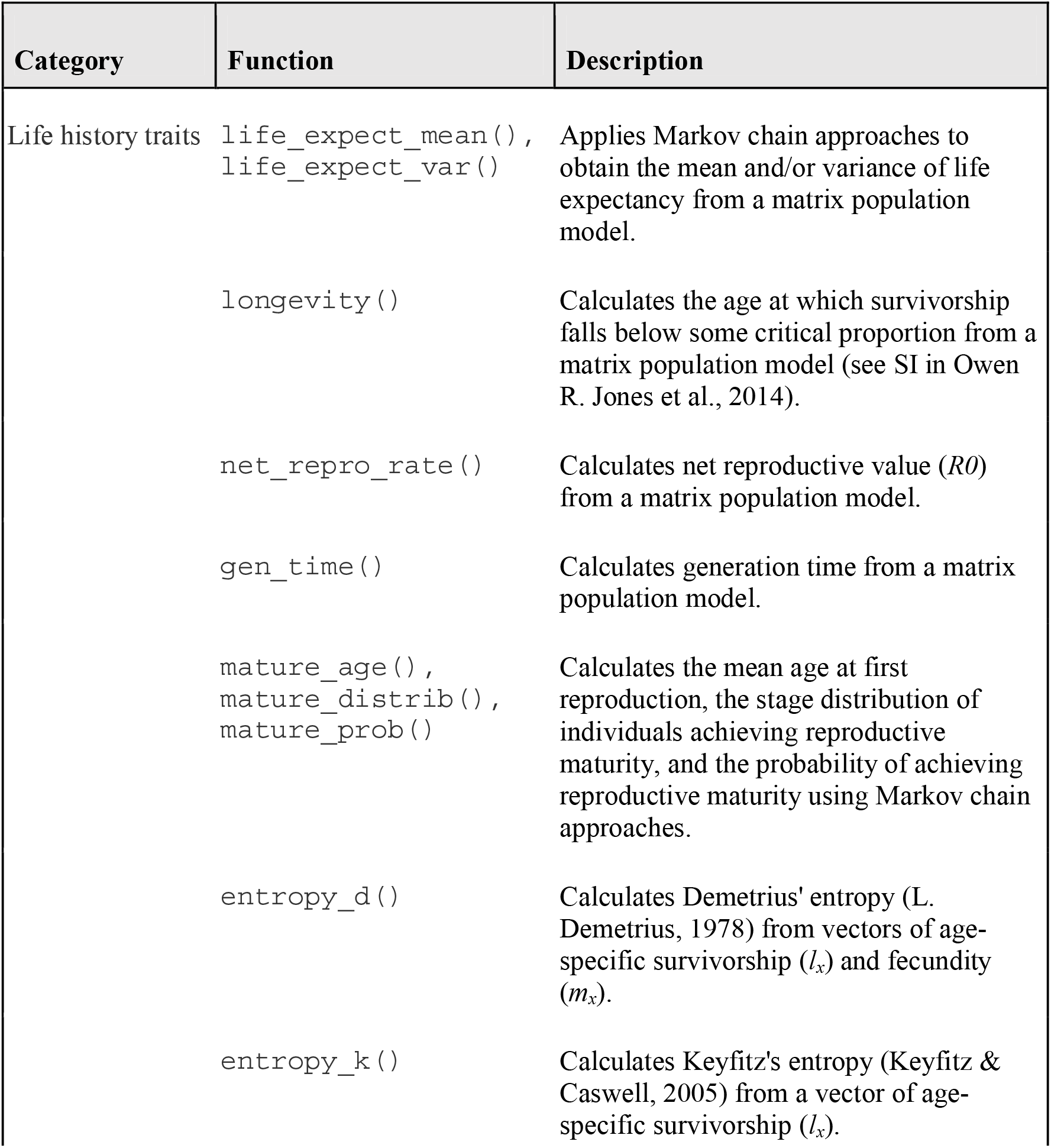
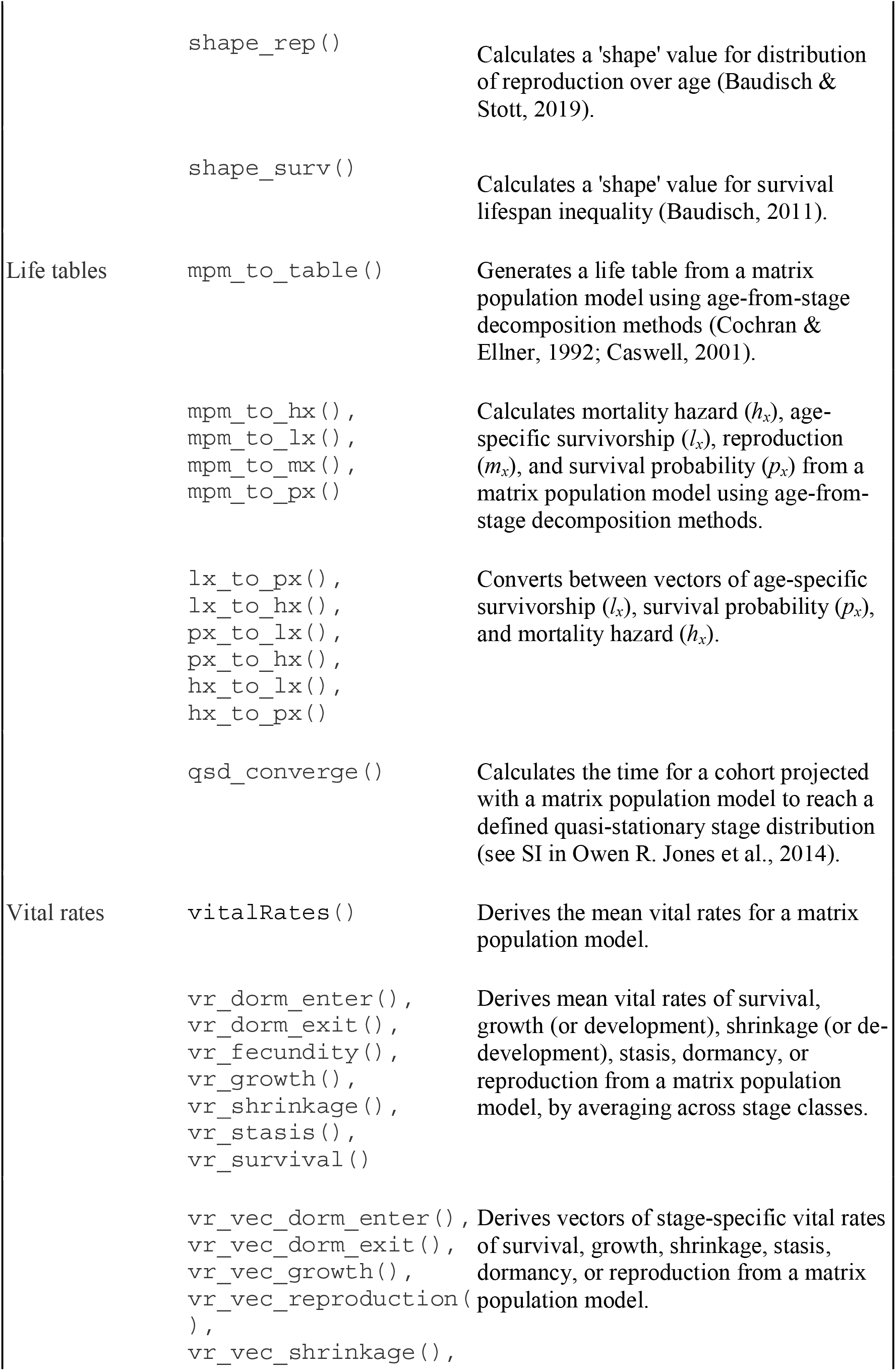
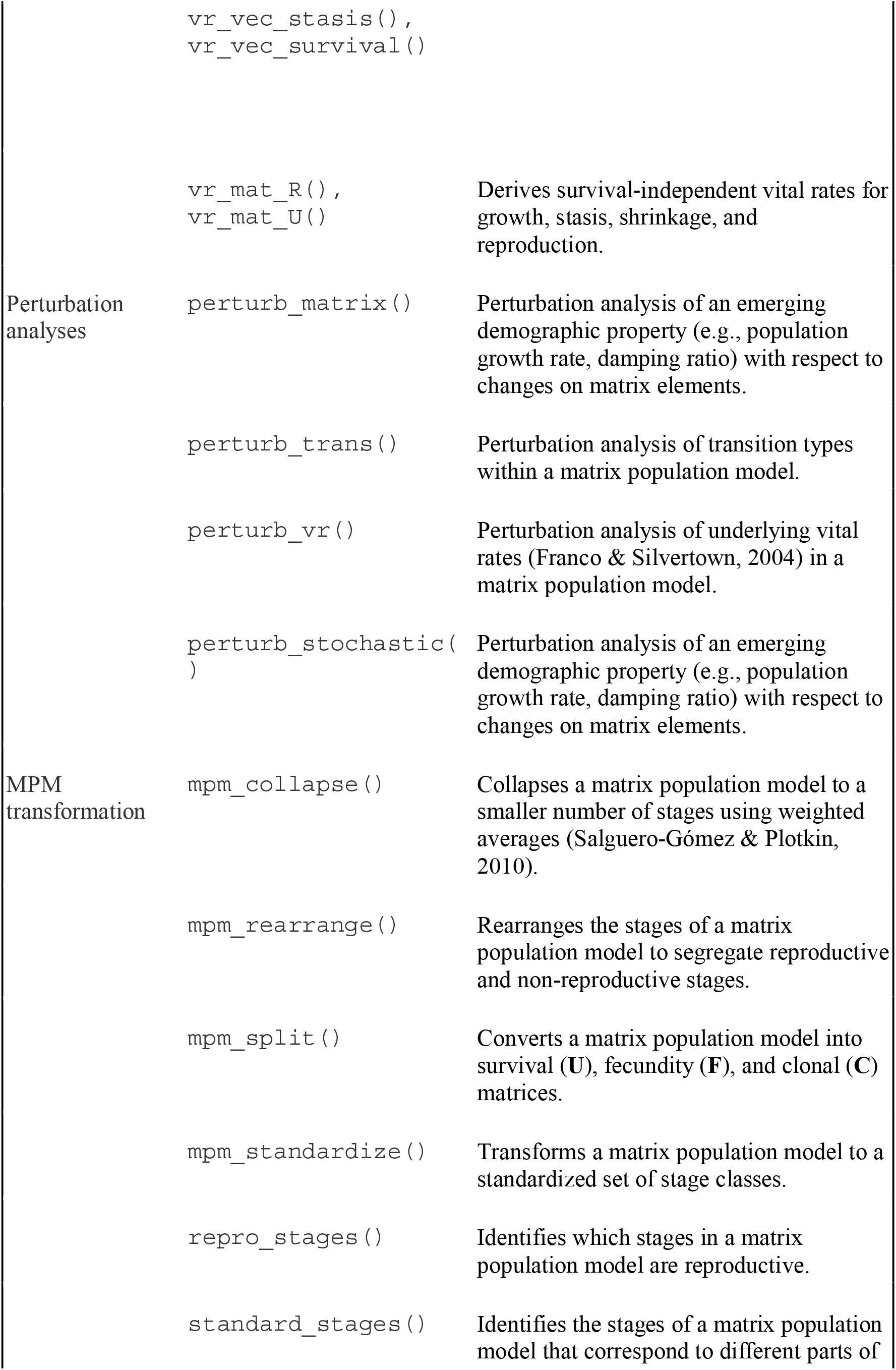
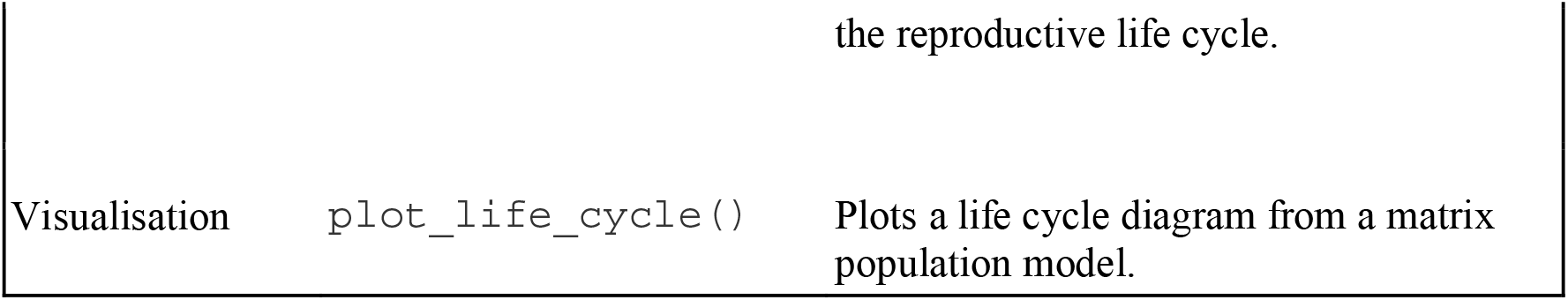
The functions in Rage are grouped into six categories: Life history traits, Life tables, Vital rates, Perturbation analyses, MPM transformation, and Visualisation. We outline the most important functions here with a brief description. Users should consult the package documentation for a full description of named functions (e.g., `?life_expect_mean`) and to see a complete list of functions.

To illustrate the functionality and inter-compatibility of functions among these categories, we describe a workflow that reconciles a common problem in comparative life history analysis: the desired life history metric requires an age-structured life table, but the available data are stage-structured MPMs. Although the mathematical descriptions for each step have long been available in the demographic literature, **`Rage`** both implements these as R functions and does so in a way that enables interoperability of function inputs and outputs. We provide in-depth vignettes for each group of functions at the **`Rage`** documentation website (https://jonesor.github.io/Rage/). However, several **`Rage`** functions, such as `mpm_to_table, entropy_…` and `shape_…`, rest on the production of age-based life tables from stage-based matrices and thus it is pertinent to outline this important aspect of **`Rage`** here.

To enable a broader range of life history analyses on data from MPMs, **`Rage`** implements conversions of stage-structured MPMs to age-specific mortality and fertility life tables using methods developed by Cochran and Ellner (1992), Caswell (2001) and Caswell et al. (2018). These methods require that MPMs are decomposed into their constituent submatrices, **U**, and optionally **F** and/or **C** (see above) and the determination of the stage we consider to be the start of the life cycle (e.g., seed establishment, seed germination, etc.). In a nutshell, the method works by an iterative procedure whereby a synthetic cohort starting at age zero is projected using the matrix model. At every iteration the cohort ages by one projection interval (often one year), and we can keep track of survivorship (*l_x_*), the proportion of the original cohort that have survived each iteration. Fecundity is calculated in an analogous way. The result is a full life table that is readily available for use in analyses that require age-, rather than stage-structured trajectories of demographic processes. We direct readers to Caswell (2001), Caswell et al. (2018) and in the supplementary information of Jones et al. (2014).

Once an *l_x_* trajectory is calculated, the other quantities of standard life tables can be calculated using standard life table calculations (Preston, Heuveline, & Guillot, 2000). In **`Rage`**, the function `mpm_to_table` applies these calculations to produce a life table that includes standard life table columns including age, survivorship, age-specific probability of death, force of mortality, remaining life expectancy. In addition, **`Rage`** provides functionality to calculate age trajectories for individual variables (i.e., subsets of the full life table) using the `mpm_to_…` set of functions (e.g., `mpm_to_lx`; Box 1).

Importantly, converting MPMs to life tables can introduce mathematical artefacts that compromise the resulting analyses. **`Rage`** provides functions to diagnose and, when possible, correct for these artefacts. All age-from-stage calculations produce age-trajectories that inevitably asymptote as a mathematical consequence of describing the vital rates as functions of discrete stages (Horvitz & Tuljapurkar, 2008). Regardless of how low the survival probabilities are in an MPM, there will be a non-zero probability that an individual could reach ages of 100, 10,000, or >1 million years. The exponential rate that these probabilities decay with increasing age is determined by the dominant eigenvalue of **U**, but even rapid decay can bias some life history metrics (e.g., entropy and life span measures). **`Rage`** provides a convenient and principled way of correcting for this artefact by imposing a lower probability threshold defined by the degree of convergence to the quasi-stationary distribution (see also the Supplementary Information of Owen R. Jones et al., 2014). In **`Rage`** we do this by first scaling the right eigenvector (**w**) so that it sums to one and then, for each iteration of the age-from-stage calculations, we measure the convergence of the proportional cohort structure as Δ_x_ = 0.5 ||**P_x_** - **w**||, where **p_x_** is the proportional stage structure at the *x*th iteration of the age-from-stage calculations (i.e., at time *x*). When **p_x_** eventually converges to equal **w**, Δ_x_ will equal 0. We can use this information to truncate the life tables produced from age-from-stage methods to, for example, ages where Δ_x_> 0.05. Furthermore, we may judge the reliability of age-from-stage methods by comparing the *l_x_* trajectory with the Δ_x_ trajectory: If convergence is reached before *l_x_* declines to, for example, 0.05 (i.e., 5% of the cohort remaining alive) we suggest reconsidering the use of this approach for that particular model.

In Box 2 we demonstrate the use of `Rage` via a global analysis of mammalian longevity introduced in Box 1. The life history metric of interest is calculated with **`Rage`**’s `longevity` function—a novel implementation in this package—by projecting a hypothetical cohort of individuals with an MPM until only a user-defined (default: 1%) fraction of individuals from the initial cohort remain alive. Since only a single cohort is tracked, the function requires only the **U** submatrix (stage-specific survival and transition rates) as the demographic process input, which may be supplied directly by the user or extracted from a `CompadreDB` object using the `matU` function from `Rcompadre`.

#### Box 2: Using Rage to calculate and visualise longevity

Here we demonstrate the use of **Rage**, focussing on the global analysis of mammalian longevity introduced in **Box 1**. We begin our mammal longevity analysis by adding columns to the data extracted from COMADRE (Box 1) that contain the two user-supplied arguments, `matU` and `start_life`, using the `dplyr` function `mutate`. We can then pair `mutate` with the base R function `mapply` to call the `longevity` function with each row’s `matU` and `start_life` arguments and return the estimated longevity in a new column. Then we check the age of convergence to the quasi-stationary stage distribution (QSD), and `filter` the data set so that it only includes matrices where the estimated longevity is less than or equal to the age at which QSD is reached. As one might expect, there is a strong association between generation time and our measure of life span (Fig. 3). It would of course be interesting to use more formal statistical methods to explore this (and similar relationships) further, for example to examine the variation in the scaling relationship across orders. When doing so it will be important to carefully consider taxonomic and geographic or ecoregion bias in the dataset. In addition, researchers should carefully vet the included data for suitability - including a consideration of whether the models are based on pre- or post-reproduction censuses.

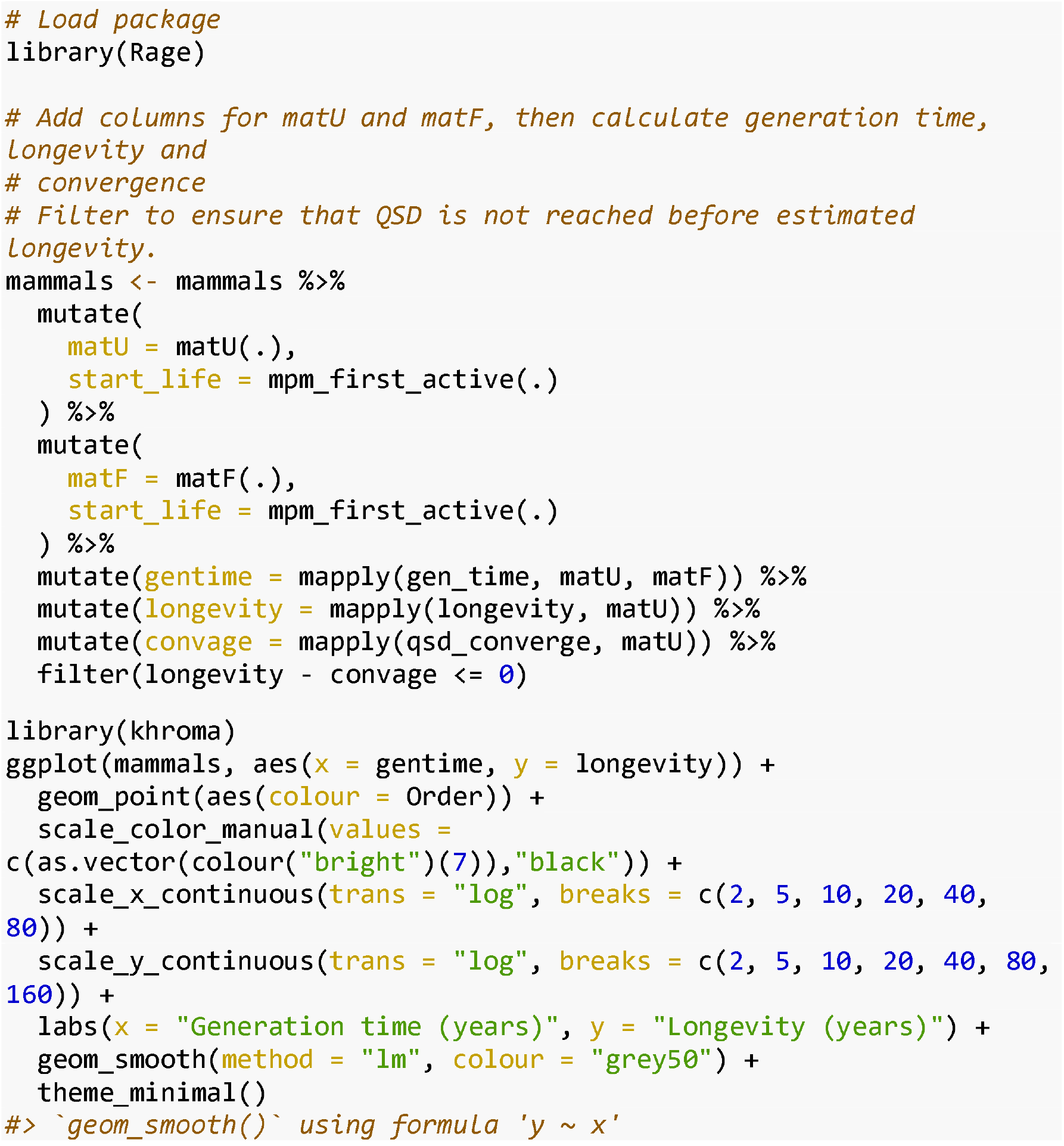

The `longevity` function also requires us to define which stage we consider to be the start of the life cycle. This is fairly clear for most mammals but may be more subjective in some groups depending on the goals of the analysis (e.g., seed maturation *vs* germination for plants with a persistent seed bank). The **`Rcompadre`** function `mpm_first_active` facilitates scaling this task across a large number of MPMs by returning an integer index for the first active stage class (i.e., non-dormant), as defined by the original study author of the MPM. Like the results of `Rcompadre::cdb_flag`, we intend this to be used as a guide—not a replacement—for careful evaluation of suitability. It may be more appropriate to identify the start of life manually in some cases. Users may control the cohort survivorship threshold via the argument `lx_crit`. The default, 0.01 (=1%) may not be suitable for all organisms, and users may find that exploring other quantiles (e.g., 50%) offers a richer description of the age-at-death distribution. Finally, the function requires us to set a maximum age to consider (`xmax`, default = 1000) as a pragmatic matter of computational speed. This default can be increased for exceptionally long-lived organisms, and we remind users that all measures of age in the **`Rage`** package use the projection interval of the MPM provided (see the `ProjectionInterval` metadata column for COM(P)ADRE data retrieved using `Rcompadre::cdb_fetch`).

## Conclusions

The tools provided by **`Rcompadre`** and **`Rage`** facilitate efficient and at-scale use of an unrivalled database of demographic process rates and the calculation of numerous life history and demographic metrics that are useful in ecology and evolution. In so doing, this pair of packages fills gaps and reduces overhead in the analytical workflow of comparative and macroecological demographic analysis. Although we designed the packages to operate together, **`Rage`** is also well-suited for general use with non-COM(P)ADRE matrix population models, whether in support of the analysis of new empirical MPMs or simulation-based theoretical studies of life history. We showcase the use of these packages to illustrate how they may be particularly useful in comparative demographic studies, for example, to address topics related to the evolution of life histories or comparative population dynamics across many species.

Users can obtain a complete index of the functions available in **`Rcompadre`** and **`Rage`** by running `?Rcompadre` and `?Rage` respectively in R, or by visiting the package documentation websites at https://jonesor.github.io/Rcompadre/ and https://jonesor.github.io/Rage/, respectively. Our ultimate hope is that democratising access to demographic data and analytic tools will empower a wide range of users to unlock the great potential of matrix population models. This will allow the community to further our basic understanding of life history, enable data-driven conservation management, and educate and inspire the next generation of population biologists.

## Supporting information

non_piped_version.pdf

## Acknowledgements

We are grateful to the Max Planck Institute for Demographic Research for funding a workshop in winter 2017 to kickstart the development of these R packages. We thank the many attendees of our teaching workshops at numerous conferences, including the annual meetings of the British Ecological Society, Evolutionary Demography Society, International Convention for Conservation Biology, and the Ecological Society of America. They inspired much of the functionality of these R packages. ORJ was supported by the Independent Research Fund Denmark (DFF-6108-00467). RS-G was supported by a NERC Independent Research Fellowship (NE/M018458/1). JC-C, ORJ, RS-G, CCT, and CS were supported by an NSF Advances in Bioinformatics Development Award (#DBI-1661342). We thank D. Buss, J. Jones, J. Metcalf, H. Caswell and B. Kendall for contributing pieces of code and/or advice at early stages of this project. We are also grateful to Y. Vindenes and two anonymous reviewers for constructive comments on an earlier draft of our manuscript.

## Authors contributions

ORJ and RS-G conceived the packages. ORJ, PB, IS, TJ, WKP, JC-C, SL, GR, CCT, CS, PC, JJ and RS-G wrote code and/or contributed to documentation. IS designed the logos and JJ and PC created Fig. 1. ORJ led the writing of the manuscript, and all authors contributed to the drafts and gave final approval to publication.

## Data availability

Data used in the examples presented here are publicly available from www.compadre-db.org.

## Supplementary materials

We provide several vignettes which guide users through most of the functionality of Rcompadre and Rage. These vignettes are available at the package development web pages at https://jonesor.github.io/Rcompadre/ and https://jonesor.github.io/Rage/, under “Articles”, in the dropdown menu.

### Rcompadre

1. Getting started with Rcompadre
2. Using Rcompadre with the tidyverse
3. Vectorising with Rcompadre
4. Obtaining references
5. Using your own matrix data

### Rage

1. Getting started with Rage
2. Deriving vital rates from an MPM
3. Deriving life history traits from an MPM
4. Age-from-stage analyses
5. Suggested quality control

An additional piece of supplementary material is a version of the code in Boxes 1 and 2 that does not use pipes: *non_piped_version.pdf*

## Footnotes

1 R includes significant support for object-oriented programming, and the S4 system is one of R’s systems for defining object classes. It is a stricter, less flexible system than R’s base system (S3) but has the advantage of enhancing consistency in how objects are defined and handled, and in the ease with which data can be accessed from nested objects. The details are far beyond the scope of this article, but see Wickham (2019) for fuller coverage.

## Notes

https://www.compadre-db.org

### Competing Interest Statement

The authors have declared no competing interest.

### Summary of Updates

This version reflects a major revision requested by reviewers. It restructures the manuscript and expands the description of our methods.

