## Supplementary material for "Rcompadre and Rage - two R packages to facilitate the use of the COMPADRE and COMADRE databases and calculation of life history traits from matrix population models": non_piped_version.pdf

### Non-piped version of Box code

The following code is a non-piped version of the code used in Boxes 1 and 2.

It requires the following packages:

```
require(tidyverse)
require(Rage)
require(Rcompadre)
require(khroma)
```

#### Box 1 - produce Figure 1.

```
comadre <- cdb_fetch("comadre", flag = TRUE)

mammals <- comadre
mammals <- filter(mammals, Class == "Mammalia")
mammals <- filter(mammals, MatrixSplit == "Divided")
mammals <- filter(
  mammals,
  check_NA_U == FALSE, check_zero_U == FALSE,
  check_zero_F == FALSE, check_zero_U_colsum == FALSE
)
mammals <- filter(mammals, ProjectionInterval == 1)

ggplot(mammals, aes(x = Lon, y = Lat)) +
  borders(database = "world", fill = "grey80", col = NA) +
  geom_point(alpha = 0.4, color = "#E69F00") +
  scale_x_continuous(breaks = seq(-180, 180, 90), expand = c(0, 0)) +
  scale_y_continuous(expand = c(0, 0)) +
  labs(x = "Longitude", y = "Latitude") +
  theme_minimal()
```

#### Box 2 - produce Figure 2.

```
mammals$matU <- matU(mammals)
mammals$matF <- matF(mammals)
mammals <- mutate(mammals, gentime = mapply(gen_time, matU, matF))
mammals <- mutate(mammals, longevity = mapply(longevity, matU))
mammals <- mutate(mammals, convage = mapply(qsd_converge, matU))
mammals <- filter(mammals, longevity - convage <= 0)
```

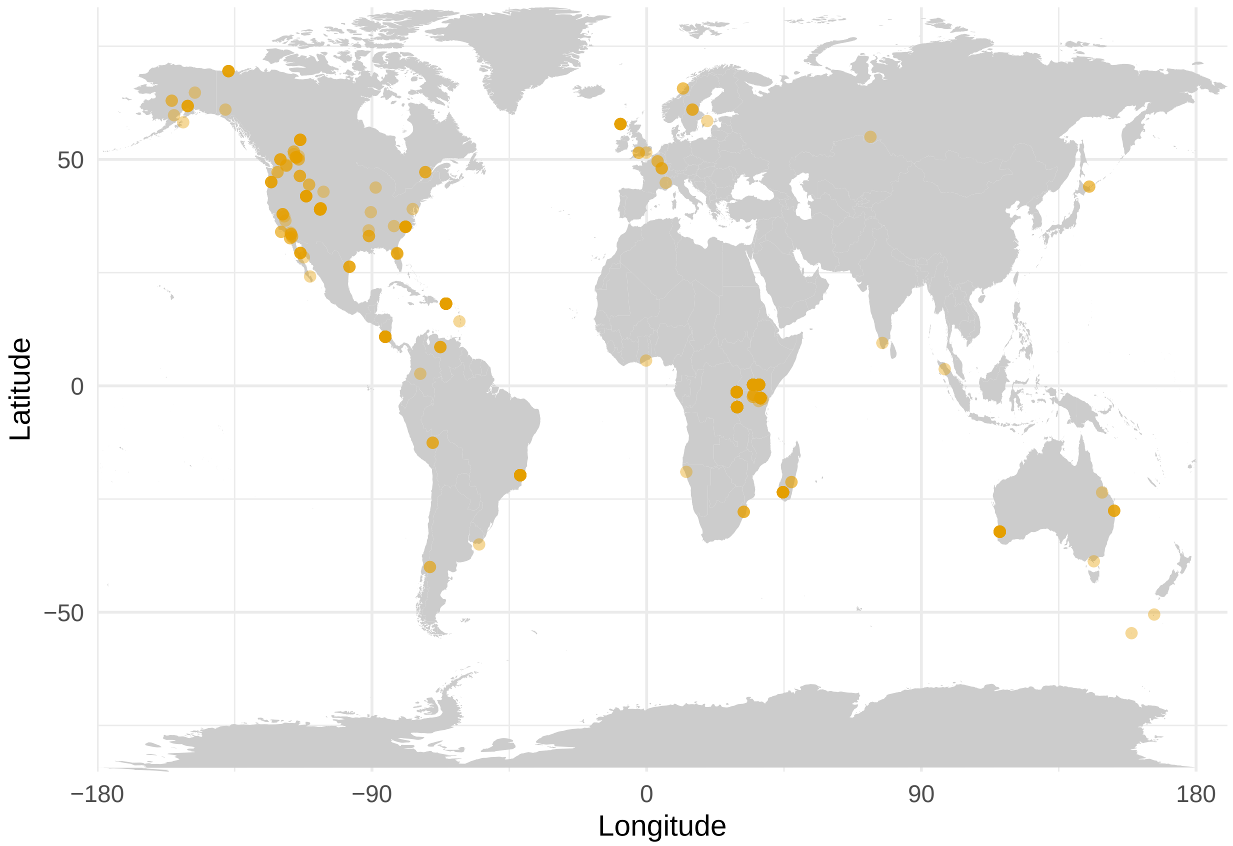


Figure 1: The geographic distribution of MPM data matching our criteria.

```
ggplot(mammals, aes(x = gentime, y = longevity)) +
  geom_point(aes(colour = Order)) +
  scale_color_manual(values = c(as.vector(colour("bright")(7)),"black")) +
  scale_x_continuous(trans = "log", breaks = c(2, 5, 10, 20, 40, 80)) +
  scale_y_continuous(trans = "log", breaks = c(2, 5, 10, 20, 40, 80, 160)) +
  labs(x = "Generation time (years)", y = "Longevity (years)") +
  geom_smooth(method = "lm", colour = "grey50") +
  theme_minimal()
```

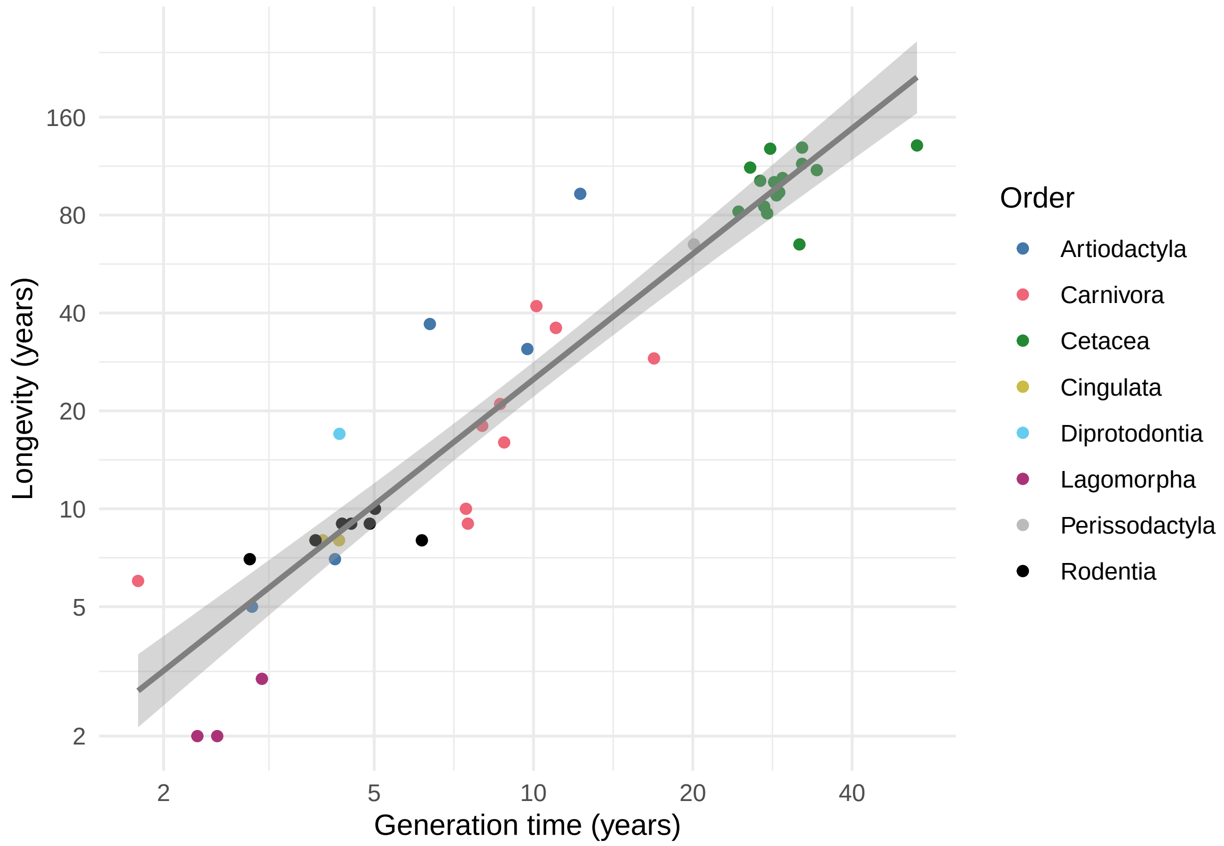


Figure 2: The relationship between generation time and longevity
